## Supplements for "Testing of the survivin suppressant YM155 in a large panel of drug-resistant neuroblastoma cell lines"

### Figure S1

## A

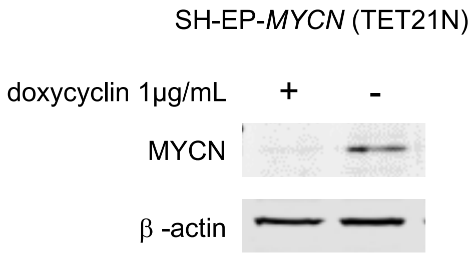


## B

| | YM155 $IC_{50}$ (nM) |
|---|---|
| - doxycyclin (1µg/mL) | 5.31 ± 3.21 |
| + doxycyclin (1µg/mL) | 4.40 ± 1.69 |

**Figure S1. Effects of MYCN expression on neuroblastoma cell sensitivity to YM155.** A) MYCN levels in SH-EP-*MYCN* (TET21N) cells in the absence or presence of doxycycline; B) YM155 concentrations that reduce the viability of SH-EP-*MYCN* (TET21N) cells by 50% ($IC_{50}$) in the absence or presence of doxycycline as determined by MTT assay after a 5-day treatment period.

#### Figure S2

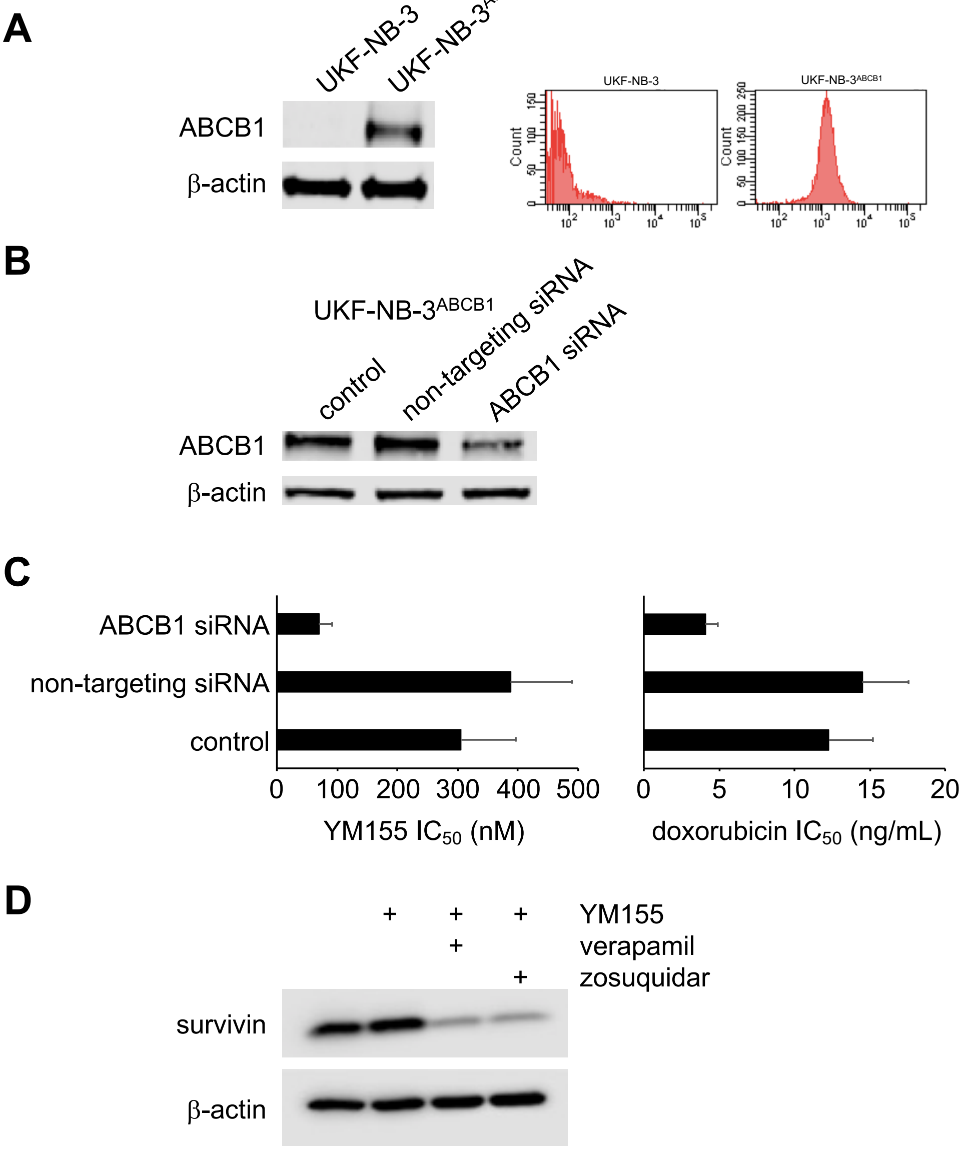


**Figure S2. Effects of YM155 in ABCB1-transduced cells.** A) Representative Western blots and flow cytometry histograms indicating ABCB1 levels in UKF-NB-3 cells and in UKF-NB-3 transduced with a lentiviral vector encoding ABCB1 (UKF-NB-$3^{ABCB1}$) . B) Effect of siRNA directed against ABCB1 on cellular ABCB1 levels in UKF-NB-$3^{ABCB1}$ cells. C) Concentrations of YM155 and doxorubicin (alternative ABCB1 substrate used as control) that reduce the viability of UKF-NB-$3^{ABCB1}$ cells by 50% ($IC_{50}$) as determined by MTT assay after 120h of incubation. D) Effects of YM155 (100nM) on survivin levels in UKF-NB-$3^{ABCB1}$ cells after 24h of incubation in the presence or absence of verapamil (5µM) or zosuquidar (1.25µM).

### Figure S3

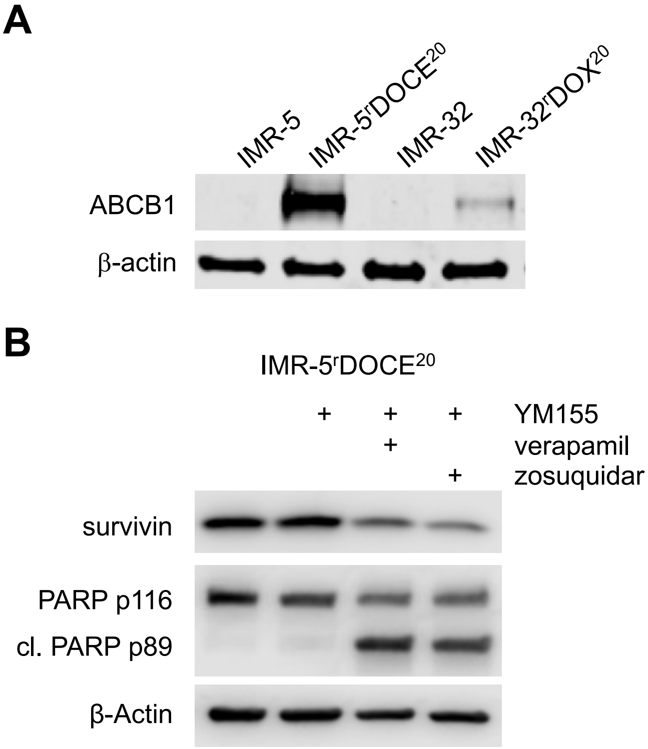


**Figure S3. ABCB1 expression and YM155 activity in drug-adapted neuroblastoma cells.** A) Representative Western blots indicating ABCB1 levels in IMR-5, IMR-5$^{r}$DOCE$^{20}$, IMR-32, and IMR-32$^{r}$DOX$^{20}$. B) Effects of YM155 (500nM) on survivin levels and PARP cleavage in IMR-5$^{r}$DOCE$^{20}$ cells in the presence or absence of verapamil (5µM) or zosuquidar (1.25µM) after 24h of incubation.

**Table S1**. YM155 concentrations that reduce the viability of neuroblastoma cell lines by 50% ($IC_{50}$) as indicated by MTT assay after 120h of incubation.

| Cell line | YM155 $IC_{50}$ (nM) | Cell line | YM155 $IC_{50}$ (nM) |
|---|---|---|---|
| Be(2)-C | 24.25 ± 2.59 | NLF$^{r}$GEMCI$^{20}$ | 1.84 ± 0.47 (0.1) |
| CHP-134 | 2.64 ± 0.50 | NLF$^{r}$IRINO$^{1000}$ | 6.93 ± 0.71 (0.3) |
| GIMEN | 33.74 ± 2.26 | NLF$^{r}$MEL$^{3000}$ | 15.36 ± 3.20 (0.6) |
| IMR-5 | 7.18 ± 1.04 | NLF$^{r}$OXALI$^{4000}$ | 33.67 ± 2.67 (1.3) |
| IMR-5$^{r}$CARBO$^{5000}$ [(1)] | 8.55 ± 2.01 (1.2)[2] | NLF$^{r}$VCR$^{10}$ | 334.5 ± 21.6 (12.5) |
| IMR-5$^{r}$CDDP$^{1000}$ | 19.71 ± 5.70 (2.7) | NLF$^{r}$VINB$^{10}$ | 38.10 ± 12.02 (1.4) |
| IMR-5$^{r}$DOCE$^{20}$ | 21549 ± 638 (3001) | NMB | 6.41 ± 1.17 |
| IMR-5$^{r}$DOX$^{20}$ | 116.3 ± 21.6 (16.2) | SHEP | 10.15 ± 0.84 |
| IMR-5$^{r}$ETO$^{100}$ | 8.29 ± 3.95 (1.2) | SHEP$^{r}$CDDP$^{1000}$ | 30.83 ± 2.24 (13.7) |
| IMR-5$^{r}$GEMCI$^{20}$ | 7.08 ± 1.20 (1.0) | SHEP$^{r}$ETO$^{100}$ | 20.24 ± 10.16 (2.0) |
| IMR-5$^{r}$MEL$^{1000}$ | 11.10 ± 1.57 (1.5) | SHEP$^{r}$VCR$^{10}$ | 20.95 ± 1.45 (2.1) |
| IMR-5$^{r}$OXALI$^{4000}$ | 10.18 ± 2.69 (1.4) | SH-SY5Y | 31.8 ± 6.50 |
| IMR-5$^{r}$TOPO$^{20}$ | 4.88 ± 1.72 (0.7) | SK-N-AS | 3.55 ± 0.21 |
| IMR-5$^{r}$VCR$^{10}$ | 472.9 ± 97.4 (65.9) | SK-N-SH | 74.94 ± 19.52 |
| IMR-5$^{r}$VINB$^{20}$ | 1608 ± 212 (224) | UKF-NB-2 | 4.18 ± 0.27 |
| IMR-5$^{r}$VINOR$^{20}$ | 4978 ± 147 (693) | UKF-NB-2$^{r}$CARBO$^{2000}$ | 318.2 ± 42.7 (76.1) |
| IMR-32 | 1.40 ± 0.35 | UKF-NB-2$^{r}$CDDP$^{1000}$ | 1.15 ± 1.21 (0.3) |
| IMR-32$^{r}$CARBO$^{1000}$ | 9.35 ± 0.97 | UKF-NB-2$^{r}$DOCE$^{10}$ | 1108 ± 179 (265) |
| IMR-32$^{r}$DOX$^{20}$ | 35.63 ± 2.23 (5.0) | UKF-NB-2$^{r}$DOX$^{20}$ | 347.0 ± 55.2 (83.0) |
| IMR-32$^{r}$ETO$^{100}$ | 1.53 ± 0.13 (0.2) | UKF-NB-2$^{r}$OXALI$^{600}$ | 3.25 ± 0.64 (0.8) |
| IMR-32$^{r}$GEMCI$^{20}$ | 2.16 ± 0.22 (0.3) | UKF-NB-2$^{r}$VCR$^{10}$ | 5940 ± 247 (1421) |
| IMR-32$^{r}$OXALI$^{800}$ | 0.60 ± 0.02 (0.1) | UKF-NB-3 | 0.49 ± 0.10 |
| IMR-32$^{r}$TOPO$^{7.5}$ | 0.45 ± 0.06 (0.1) | UKF-NB-3$^{r}$CARBO$^{2000}$ | 155.4 ± 24.6 (317) |
| IMR-32$^{r}$VINOR$^{5}$ | 16.43 ± 1.08 (2.3) | UKF-NB-3$^{r}$CDDP$^{1000}$ | 5.32 ± 1.21 (10.9) |
| LAN-6 | 248.1 ± 32.9 | UKF-NB-3$^{r}$DOCE$^{10}$ | 469.6 ± 113.1 (958) |
| NB-S-124 | 76.66 ± 6.51 | UKF-NB-3$^{r}$DOX$^{20}$ | 15,700 ± 1,019 (32041) |
| NGP | 12.48 ± 3.01 | UKF-NB-3$^{r}$ETO$^{200}$ | 7.97 ± 0.13 (16.3) |
| NGP$^{r}$CARBO$^{5000}$ | 112.3 ± 5.0 (9.0) | UKF-NB-3$^{r}$GEMCI$^{10}$ | 0.40 ± 0.01 (0.8) |
| NGP$^{r}$CDDP$^{1000}$ | 13.00 ± 0.42 (1.0) | UKF-NB-3$^{r}$Nutlin$^{10 \mu M}$ | 1.18 ± 0.07 (2.4) |
| NGP$^{r}$DACARB$^{18}$ | 20.59 ± 1.84 (1.6) | UKF-NB-3$^{r}$OXALI$^{4000}$ | 1.80 ± 0.78 (3.7) |
| NGP$^{r}$DOCE$^{20}$ | 159.0 ± 19.5 (12.7) | UKF-NB-3$^{r}$TOPO$^{20}$ | 7.40 ± 0.71 (15.1) |
| NGP$^{r}$DOX$^{20}$ | 306.9 ± 78.5 (24.6) | UKF-NB-3$^{r}$VCR$^{10}$ | 26.59 ± 6.37 (54.3) |
| NGP$^{r}$ETO$^{400}$ | 59.20 ± 11.40 (4.7) | UKF-NB-6 | 0.65 ± 0.09 |
| NGP$^{r}$GEMCI$^{20}$ | 41.55 ± 6.13 (3.3) | UKF-NB-6$^{r}$CARBO$^{2000}$ | 16.83 ± 1.62 (25.9) |
| NGP$^{r}$MEL$^{3000}$ | 26.10 ± 3.86 (2.1) | UKF-NB-6$^{r}$CDDP$^{2000}$ | 79.93 ± 7.14 (123) |
| NGP$^{r}$OXALI$^{4000}$ | 6.93 ± 0.28 (0.6) | UKF-NB-6$^{r}$DOCE$^{10}$ | 14.33 ± 4.08 (22.0) |
| NGP$^{r}$VCR$^{20}$ | 6986 ± 715 (560) | UKF-NB-6$^{r}$DOX$^{20}$ | 11.80 ± 1.56 (18.2) |
| NLF | 26.78 ± 4.04 | UKF-NB-6$^{r}$ETO$^{200}$ | 3.60 ± 0.01 (5.5) |
| NLF$^{r}$CARBO$^{5000}$ | 340.5 ± 34.5 (12.7) | UKF-NB-6$^{r}$GEMCI$^{10}$ | 2.10 ± 0.84 (3.2) |
| NLF$^{r}$CDDP$^{500}$ | 12.58 ± 5.39 (0.5) | UKF-NB-6$^{r}$OXALI$^{4000}$ | 5.34 ± 0.71 (8.2) |
| NLF$^{r}$DOCE$^{20}$ | 21.6 ± 5.98 (0.8) | UKF-NB-6$^{r}$TOPO$^{20}$ | 3.47 ± 0.81 (5.3) |
| NLF$^{r}$DOX$^{40}$ | 34.88 ± 4.33 (1.3) | UKF-NB-6$^{r}$VCR$^{10}$ | 49.30 ± 2.24 (75.8) |
| NLF$^{r}$ETO$^{100}$ | 7.40 ± 0.54 (0.3) | UKF-NB-6$^{r}$VINOR$^{40}$ | 228.5 ± 41.5 (352) |

[1] CARBO, carboplatin; CDDP, cisplatin; DACARB, dacarbazine; DOX, doxorubicin; ETO, etoposide; GEMCI, gemcitabine; IRINO, irinotecan; MEL, melphalan; Nutlin, nutlin-3; OXALI, oxaliplatin; TOPO, topotecan; VCR, vincristine; VINB, vinblastine; VINOR, vinorelbine

**Table S2**. YM155 concentrations that reduce the viability of MYCN-amplified and non-MYCN-amplified neuroblastoma cell lines by 50% (IC50) in the absence or presence of the ABCB1 inhibitors verapamil (5 µM) or zosuquidar (1.25 µM) as indicated by MTT assay after 120h of incubation.

| **Cell line** | **YM155 $IC_{50}$ (nM)** | **+ verapamil (5 µM)[1] YM155 $IC_{50}$ (nM)** | **+ zosuquidar (1.25 µM) YM155 $IC_{50}$ (nM)** |
|---|---|---|---|
| *MYCN amplification* | | | |
| CHP-134 (wt)[2] | 2.64 ± 0.50 | 1.64 ± 0.27 | 1.85 ± 0.34 |
| IMR-5 (wt) | 7.18 ± 1.04 | 9.70 ± 1.97 | 10.64 ± 2.80 |
| IMR-32 (wt) | 1.40 ± 0.35 | 1.70 ± 0.41 | 1.80 ± 0.23 |
| NB-S-124 (wt) | 76.66 ± 6.51 | 12.52 ± 1.16 | 3.20 ± 0.40 |
| NGP (wt) | 12.48 ± 3.01 | 17.35 ± 4.97 | 24.95 ± 0.21 |
| NLF (V203M) | 26.78 ± 4.04 | 19.55 ± 1.20 | 45.30 ± 1.34 |
| UKF-NB-2 (wt) | 4.18 ± 0.27 | 4.55 ± 0.32 | 2.85 ± 0.14 |
| UKF-NB-3 (wt) | 0.49 ± 0.10 | 0.61 ± 0.13 | 0.74 ± 0.10 |
| UKF-NB-6 (wt) | 0.65 ± 0.09 | 0.58 ± 0.07 | 0.57 ± 0.07 |
| *no MYCN amplification* | | | |
| GIMEN (wt) | 33.74 ± 2.26 | 52.90 ± 8.62 | 50.87 ± 5.91 |
| LAN-6 (wt) | 248.1 ± 32.9 | 46.75 ± 2.33 | 24.35 ± 1.06 |
| SHEP (wt) | 10.15 ± 0.84 | 3.92 ± 0.11 | 3.20 ± 0.14 |
| SK-N-AS (null) | 3.55 ± 0.21 | 1.01 ± 0.26 | 1.31 ± 0.11 |
| SK-N-SH (wt) | 74.94 ± 19.52 | 6.80 ± 0.83 | 1.72 ± 0.15 |

[1] Effects of verapamil or zosuquidar alone on cell viability are presented in Table S5.
[2] *TP53* status: wt, wild-type; otherwise type of mutation is provided

**Table S3**. YM155 concentrations that reduce the viability of neuroblastoma cell lines with varying p53 status by 50% ($IC_{50}$) as indicated by MTT assay after 120h of incubation.

| **Cell line** | **p53 status** | **YM155 $IC_{50}$ (nM)** |
|---|---|---|
| UKF-NB-3 | wild-type | 0.49 ± 0.10 |
| UKF-NB-3$^{r}$Nutlin$^{10\mu M}$ | G245C (homo)[1] | 1.18 ± 0.07 (2.4)[2] |
| UKF-NB-3clone1 | wild-type | 0.35 ± 0.07 |
| UKF-NB-3clone1$^{r}$Nutlin$^{10\mu M}$I | stop codon in exon 4 | 0.40 ± 0.12 (1.1) |
| UKF-NB-3clone1$^{r}$Nutlin$^{10\mu M}$III | R248W (het) | 0.60 ± 0.08 (1.7) |
| UKF-NB-3clone1$^{r}$Nutlin$^{10\mu M}$IV | V173L (het) | 0.45 ± 0.06 (1.3) |
| UKF-NB-3clone1$^{r}$Nutlin$^{10\mu M}$VI | R196Q (het) | 0.55 ± 0.17 (1.6) |
| UKF-NB-3clone1$^{r}$Nutlin$^{10\mu M}$VIII | Y236C (het) | 0.50 ± 0.14 (1.4) |
| UKF-NB-3clone1$^{r}$Nutlin$^{10\mu M}$X | P151R (het) | 0.73 ± 0.08 (2.1) |
| UKF-NB-3clone3 | wild-type | 0.45 ± 0.06 |
| UKF-NB-3clone3$^{r}$Nutlin$^{10\mu M}$I | P152L (het) | 1.50 ± 0.06 (3.3) |
| UKF-NB-3clone3$^{r}$Nutlin$^{10\mu M}$VIII | N239S (het) | 0.50 ± 0.08 (1.1) |
| UKF-NB-3clone3$^{r}$Nutlin$^{10\mu M}$IX | R280S (het) | 1.03 ± 0.03 (2.3) |
| UKF-NB-3clone3$^{r}$Nutlin$^{10\mu M}$X | I251F (het) | 0.58 ± 0.09 (1.3) |
| UKF-NB-6 | wild-type | 0.65 ± 0.09 |
| UKF-NB-6$^{r}$Nutlin$^{10\mu M}$ | K132N (het); P223L (hom) | 0.64 ± 0.04 (1.0) |
| UKF-NB-6$^{r}$Nutlin$^{10\mu M}$I | S241F (hom) | 0.57 ± 0.01 (0.9) |
| UKF-NB-6$^{r}$Nutlin$^{10\mu M}$IV | C135F (het); D281Y (het) | 0.43 ± 0.04 (0.7) |

[1] homo = homozygous, het = heterozygous

[2] fold change YM155 $IC_{50}$ nutlin-3-resistant sub-line/ YM155 $IC_{50}$ respective parental cell line

**Table S4**. Mean YM155 concentrations that reduce the viability of neuroblastoma cell lines with resistance to certain drug classes by 50% ($IC_{50}$) as indicated by MTT assay after 120h of incubation. Values are presented as mean ± S.D. Individual values are presented in Table 1.

| **Drug class** | **YM155 $IC_{50}$ (nM)** |
|---|---|
| topoisomerase I inhibitors | 4.63 ± 2.52 |
| parental | 7.91 ± 8.27 |
| nucleoside analogue (gemcitabine) | 9.18 ± 14.62 |
| alkylating agents | 18.29 ± 5.63 |
| platinum drugs | 58.43 ± 98.80 |
| topoisomerase II inhibitors | 73.67 ± 112.42 |
| including UKF-NB-3$^r$DOX[20] | 1190 ± 4026 |
| taxane (docetaxel) | 354 ± 411 |
| including IMR-5$^r$DOCE[20] | 3889 ± 7908 |
| vinca alkaloids | 1725 ± 2519 |

**Table S5**. YM155 concentrations that reduce the viability of neuroblastoma cell lines by 50% (IC50) in the absence or presence of the ABCB1 inhibitors verapamil (5 µM) or zosuquidar (1.25 µM) as indicated by MTT assay after 120h of incubation.

| | | **+ verapamil (5 µM)** | | **+ zosuquidar (1.25 µM)** | |
|---|---|---|---|---|---|
| **Cell line** | **YM155 $IC_{50}$ (nM)** | **verapamil alone** | **YM155 $IC_{50}$ (nM)** | **zosuquidar alone** | **YM155 $IC_{50}$ (nM)** |
| CHP-134 | 2.64 ± 0.50 | 94 ± 13[1] | 1.64 ± 0.27 (1.6)[2] | 105 ± 6[1] | 1.85 ± 0.34 (2.4) |
| GIMEN | 33.74 ± 2.26 | 105 ± 3 | 52.90 ± 8.62 (0.6) | 92 ± 7 | 50.87 ± 5.91 (0.7) |
| IMR-5 | 7.18 ± 1.04 | 109 ± 8 | 9.70 ± 1.97 (0.7) | 104 ± 11 | 10.64 ± 2.80 (0.7) |
| IMR-5$^r$CARBO$^{5000}$ | 8.55 ± 2.01 | 91 ± 16 | 7.80 ± 0.28 (1.1) | 105 ± 8 | 27.01 ± 3.04 (0.3) |
| IMR-5$^r$CDDP$^{1000}$ | 19.71 ± 5.70 | 88 ± 11 | 15.23 ± 4.21 (1.3) | 100 ± 6 | 33.47 ± 6.84 (0.6) |
| IMR-5$^r$DOCE$^{20}$ | 21549 ± 638 | 90 ± 7 | 149.01 ± 1.99 (145) | 112 ± 2 | 13.63 ± 5.54 (1581) |
| IMR-5$^r$DOX$^{20}$ | 116.3 ± 21.6 | 97 ± 9 | 17.60 ± 0.57 (6.6) | 99 ± 8 | 13.45 ± 2.45 (8.6) |
| IMR-5$^r$ETO$^{100}$ | 8.29 ± 3.95 | 95 ± 10 | 6.99 ± 2.79 (1.2) | 97 ± 4 | 18.26 ± 3.19 (0.5) |
| IMR-5$^r$GEMCI$^{20}$ | 7.08 ± 1.20 | 108 ± 6 | 7.90 ± 2.09 (0.9) | 105 ± 8 | 12.73 ± 3.34 (0.6) |
| IMR-5$^r$MEL$^{1000}$ | 11.10 ± 1.57 | 92 ± 8 | 6.63 ± 1.30 (1.7) | 107 ± 6 | 12.80 ± 1.22 (0.9) |
| IMR-5$^r$OXALI$^{4000}$ | 10.18 ± 2.69 | 96 ± 8 | 15.80 ± 1.77 (0.6) | 110 ± 6 | 16.81 ± 2.71 (0.6) |
| IMR-5$^r$TOPO$^{20}$ | 4.88 ± 1.72 | 100 ± 13 | 5.94 ± 1.31 (0.8) | 101 ± 7 | 11.77 ± 3.95 (0.4) |
| IMR-5$^r$VCR$^{10}$ | 472.9 ± 97.4 | 93 ± 8 | 13.05 ± 2.90 (36) | 94 ± 5 | 19.35 ± 0.07 (24) |
| IMR-5$^r$VINB$^{20}$ | 1608 ± 212 | 93 ± 7 | 9.34 ± 0.94 (172) | 93 ± 6 | 10.05 ± 1.06 (160) |
| IMR-32 | 1.40 ± 0.35 | 102 ± 7 | 1.70 ± 0.41 (0.8) | 101 ± 3 | 1.80 ± 0.23 (0.8) |
| IMR-32$^r$DOX$^{20}$ | 35.63 ± 2.23 | 92 ± 15 | 1.75 ± 0.77 (20) | 89 ± 13 | 0.94 ± 0.08 (38) |
| IMR-32$^r$ETO$^{100}$ | 1.53 ± 0.13 | 90 ± 5 | 1.60 ± 0.27 (1.0) | 104 ± 6 | 3.55 ± 0.21 (2.2) |
| IMR-32$^r$GEMCI$^{20}$ | 2.16 ± 0.22 | 105 ± 8 | 1.15 ± 0.05 (1.9) | 107 ± 11 | 4.20 ± 0.45 (0.5) |
| IMR-32$^r$OXALI$^{800}$ | 0.60 ± 0.02 | 99 ± 8 | 0.71 ± 0.08 (0.8) | 105 ± 12 | 1.18 ± 0.07 (1.7) |
| IMR-32$^r$TOPO$^{7.5}$ | 0.45 ± 0.06 | 97 ± 10 | 0.61 ± 0.07 (0.7) | 101 ± 2 | 0.97 ± 0.04 (0.5) |
| LAN-6 | 248.1 ± 32.9 | 99 ± 8 | 46.75 ± 2.33 (5.3) | 103 ± 5 | 24.35 ± 1.06 (10.2) |
| NB-S-124 | 76.66 ± 6.51 | 103 ± 6 | 12.52 ± 1.16 (6.1) | 110 ± 8 | 3.20 ± 0.40 (24.0) |
| NGP | 12.48 ± 3.01 | 91 ± 8 | 17.35 ± 4.97 (0.7) | 109 ± 2 | 24.95 ± 0.21 (0.5) |
| NGP$^r$CARBO$^{5000}$ | 112.3 ± 5.0 | 112 ± 9 | 76.10 ± 3.17 (1.5) | 107 ± 5 | 158.24 ± 9.34 (0.7) |
| NGP$^r$CDDP$^{1000}$ | 13.00 ± 0.42 | 104 ± 5 | 19.61 ± 1.35 (0.7) | 101 ± 18 | 17.80 ± 0.97 (0.7) |
| NGP$^r$DACARB$^{18}$ | 20.59 ± 1.84 | 107 ± 2 | 26.26 ± 4.77 (0.8) | 103 ± 7 | 41.90 ± 5.27 (0.5) |
| NGP$^r$DOX$^{20}$ | 306.9 ± 78.5 | 92 ± 6 | 5.52 ± 0.35 (56) | 90 ± 2 | 0.70 ± 0.04 (438) |
| NGP$^r$ETO$^{400}$ | 59.20 ± 11.40 | 98 ± 16 | 50.14 ± 16.45 (1.2) | 98 ± 3 | 39.12 ± 7.87 (1.5) |
| NGP$^r$GEMCI$^{20}$ | 41.55 ± 6.13 | 94 ± 13 | 73.43 ± 16.41 (0.6) | 105 ± 12 | 10.50 ± 1.34 (4.0) |
| NGP$^r$MEL$^{3000}$ | 26.10 ± 3.86 | 99 ± 10 | 24.34 ± 1.76 (1.1) | 108 ± 9 | 18.75 ± 4.64 (1.4) |
| NGP$^r$OXALI$^{4000}$ | 6.93 ± 0.28 | 102 ± 8 | 12.25 ± 2.78 (0.6) | 101 ± 12 | 8.21 ± 1.04 (0.8) |
| NGP$^r$VCR$^{20}$ | 6986 ± 715 | 100 ± 10 | 157.60 ± 11.79 (44) | 106 ± 15 | 16.20 ± 1.74 (431) |
| NLF | 4.18 ± 0.27 | 93 ± 8 | 4.55 ± 0.32 (0.9) | 99 ± 5 | 2.85 ± 0.14 (1.5) |

**Table S6**. YM155 concentrations that reduce the viability of neuroblastoma cell lines by 50% ($IC_{50}$) in the absence or presence of zosuquidar (1.25µM) as indicated by MTT assay or CellTiterGlo after 120h of incubation.

| **Cell line** | **YM155 $IC_{50}$ (nM)** | **+ Zosuquidar[1]** |
|---|---|---|
| IMR-5 (MTT) | 7.18 ± 1.04 | 10.64 ± 2.80 |
| IMR-5 (CellTiterGlo) | 6.27 ± 1.56 | 19.99 ± 5.97 |
| IMR-5$^{r}$DOCE$^{20}$ (MTT) | 21,549 ± 638 | 13.63 ± 5.54 |
| IMR-5$^{r}$DOCE$^{20}$ (CellTiterGlo) | 32,946 ± 4360 | 14.82 ± 4.66 |
| IMR-32 (MTT) | 1.40 ± 0.35 | 1.80 ± 0.23 |
| IMR-32 (CellTiterGlo) | 1.22 ± 0.21 | 1.91 ± 1.16 |
| IMR-32$^{r}$DOX$^{20}$ (MTT) | 35.53 ± 2.23 | 0.94 ± 0.08 |
| IMR-32$^{r}$DOX$^{20}$ (CellTiterGlo) | 29.20 ± 3.56 | 0.86 ± 0.06 |

[1] The effects of zosuquidar alone are provided in Table S5.

**Table S7**. YM155 concentrations that reduce the viability of neuroblastoma cell lines by 50% ($IC_{50}$) in the absence or presence of the ABCC1 inhibitor MK571 (10µM) as indicated by MTT assay after 120h of incubation.

| | | + MK571 | |
|---|---|---|---|
| **Cell line** | **YM155 $IC_{50}$ (nM)** | **MK571 alone[1]** | **YM155 $IC_{50}$ (nM)** |
| NLF | 26.3 ± 5.9 | 98 ± 15 | 25.6 ± 9.1 |
| NLF$^r$VCR$^{10}$ | 324 ± 79 | 102 ± 17 | 141 ± 38 |

[1] Effect of MK571 (10µM) on cell viability in percentage relative to untreated control.
